# Aberrant expression of collagen family genes in the brain regions developing under agonistic interactions in male mice

**DOI:** 10.1101/276063

**Authors:** D.A. Smagin, A.G. Galyamina, I.L. Kovalenko, V.N. Babenko, N.N. Kudryavtseva

**Affiliations:** Laboratory of Neuropathology Modeling, Institute of Cytology and Genetics, Siberian Branch of Russian Academy of Sciences, Novosibirsk, Russia

**Keywords:** collagen genes, social defeats, depression, repeated aggression, extracellular matrix, brain regions

## Abstract

As previously established, chronic agonistic interactions lead to the development of depression-like state under social defeat stress in the defeated mice and pathology of aggressive behavior in the winning mice. According to the numerous research data, these psychopathological states are accompanied by tremendous molecular and cellular changes in the brain. The paper aimed to study the influence of 20-day period of agonistic interactions on the expression mode of collagen family genes, encoding the proteins, which are basic components of extracellular matrix (ECM), in the different brain regions of mice using the RNA-Seq database. Most of the differentially expressed collagen genes were upregulated in the hypothalamus and striatum of chronically aggressive and defeated mice and in the hippocampus of the defeated mice. In the ventral tegmental area the most genes were downregulated in both experimental groups. It has been assumed that aberrant expression of collagen genes induced by long experience of agonistic interactions can indicate defects of ECM specific for brain regions in mice with alternative social experiences. This study first shows remodeling of molecular base in the ECM under development of experimental psychoneuropathologies.

Corresponding authors: Kudryavtseva N.N.; Babenko V.N.

## Introduction

According to the research data, chronic agonistic interactions lead to the development of behavioral psychopathologies. They include the development of mixed anxiety/depression-like states in defeated male mice [Kudryavtseva *et al.* 1991; Kudryavtseva & Avgustinovich, 1998; Berton *et al.* 2006; Bondar et al. 2009; Galyamina *et al.* 2017], similar to those in humans, and psychosis-like behavior in the repeatedly aggressive mice [Kudryavtseva 2006; Kudryavtseva *et al.* 2014; Ibrahim *et al.* 2016]. After daily agonistic interactions, the behaviors of both social groups are crucially changed. Depressive males demonstrated total behavioral deficit (low locomotor and exploratory activity, immobility in different experimental situations), avoidance of social contacts and reduced communication, anhedonia and generalized anxiety. Chronically aggressive mice displayed enhanced aggression, high impulsivity and anxiety, as well as impaired social recognition, hyperactivity and stereotypic behaviors. Aggressive motivation prevailed in all social interactions.

It was established that under chronic social defeat stress the adult brain undergoes numerous molecular and cellular changes [Kudryavtseva *et al.* 2004; Berton *et al.* 2006; Boyarskikh *et al.* 2013]. Analyzing the whole transcriptome in brain of depressive mice, we have found changes in the expression of serotonergic [Smagin *et al.* 2013; Kudryavtseva *et al.* 2017], dopaminergic [Kovalenko *et al.* 2016], glutamatergic and GABAergic [Smagin *et al.* 2017; Galyamina *et al.* 2017] genes as well as ribosomal and mitoribosomal genes [Smagin *et al.* 2016a; Smagin *et al.* 2016b] in different brain regions. Changes in DNA methylation, histone acetylation and chromatin remodeling often accompany these changes [Hollis *et al.* 2010; Kenworthy *et al.* 2014], as well as decreases in hippocampal neurogenesis rates [Ferragud *et al.* 2010; Lagace *et al.* 2010; Van Bokhoven *et al.* 2011] in defeated mice. We found changes in activity of monoaminergic gene expression in brain regions, activation of cell proliferations in the hippocampus, as well as growth of new neurons under repeated experience of aggression in male mice [Kudryavtseva *et al.* 2004; Bondar *et al.* 2009; Filipenko *et al.* 2001; Smagin *et al.* 2013; Smagin *et al.* 2015; Kudryavtseva *et al.* 2017]. Using the transcriptome analysis on RNA-seq data this report is aimed to study the changes of the collagen family gene expression in the 5 brain regions of the male mice with alternative social experience in daily agonistic interactions over a 20-day period.

There are 28 collagen subfamilies (Col1 – Col28) comprising 45 collagen genes total in mouse genome [Myllyharju & Kivirikko 2004; Gordon *et al.* 2010; Ricard-Blum, 2011; Seppänen e*t al*, 2006, Seppänen *et al*, 2007]. The collagen family genes (*Col*\*) encode collagen proteins which manifest the predominant glycoproteins group of extracellular matrix (ECM) constituting the main component of connective tissue in mammals [Myllyharju & Kivirikko 2004; Gordon *et al* 2010; Ricard-Blum 2011]. Collagen is a fibrous protein found in vertebrates representing the major element of skin, bone, tendon, cartilage, blood vessels and teeth. The role of collagens is forming fibrils and assembling into elongated fibers in the ECM [Fox 2008].

In the brain tissue a fibrillar collagen is apparently absent [Hubert *et al*. 2009], while the collagens function as structural and active adhesion molecules [Myllyharju & Kivirikko 2004; Kadler *et al*. 2007]. Being the structural components of extracellular matrix, collagens are also involved in neuronal development of the brain and play a role in regulation of axonal outgrowth, and synaptic differentiation [Sertie, *et al*. 2000; Schneider & Granato 2006; Fox *et al*. 2007], neural maturation [Fox *et al*. 2007; Heffron *et al*. 2009], synaptogenesis and establishment of brain architecture [Hubert *et al*. 2009; Chernousov *et al*. 2006]. Some of the collagen proteins are expressed by neurons [Claudepierre *et al*. 2005; Hashimoto *et al*. 2002; Sund *et al*. 2001].

Most CNS pathologies associated with collagens are related to neurodevelopment. For example, it was shown that collagen type IV promotes the differentiation of neuronal progenitors and inhibits astroglial differentiation in cortical cell cultures [Ali *et al*. 1998]. Post-mortem human studies showed that collagen type XVII is widely expressed in the brain and is located primarily in the soma and proximal axons of neurons in contrast to glial cells, which do not express this collagen [Seppänen *et al*. 2007]. Collagen XVI may act as a substrate for adhesion and invasion of connective tissue tumor cells. Alteration of tissue location and expression level appears to promote tumorigenesis and inflammatory reactions [Grässel & Bauer 2013]. Thus, altering the cell-matrix interaction through collagen XVI might be a molecular mechanism to further augment the invasive phenotype of glioma cells. It is supposed, that collagen XVI plays a decisive role in the interaction of connective tissue cells with their ECM, which is impaired in pathological situations. Amyloid-beta peptides, widely presumed to cause Alzheimer’s disease, increased mouse neuronal expression of collagen VI [Cheng *et al.* 2009]. Authors discovered that the cellular source of the collagen VI is neurons. Collagen IV proteins were increased in Alzheimer’s disease patients in frontal and temporal cortex. Significant ECM changes occur during the early stages of this disease [Lepelletier *et al.* 2017]. Col25a1 alleles have been associated with increased risk for Alzheimer’s disease [Forsell *et al.* 2010]. In these patients extracellular collagenous Alzheimer amyloid plaque component [Hashimoto *et al.* 2002], which is extracellular part of the transmembrane collagen XXV, is expressed by neurons. Interestingly, over-expression of the *Col25a1* gene in neurons in transgenic mice leads to Alzheimer’s disease-like brain pathology [Tong *et al.* 2010]. It has been supposed that enhanced expression of the ECM related with cell adhesion genes, *Col8a1* in particular, in the prefrontal cortex may affect cortical neural plasticity, including morphological neuronal changes and the afferent and/or efferent neural pathways participating in stress-related emotional behavioral patterns [Li *et al.* 2013]. All these data indicate the possible role of *Col\** family genes in different neurological disorders, however little known about reaction of brain *Col\** genes expression on chronic functional uploads and in psychoemotional disorders.

## Materials and Methods

### Animals

Adult male mice C57BL/6J was obtained from Animal Breeding Facility, Branch of Institute of Bioorganic Chemistry of the RAS (ABF BIBCh, RAS), Pushchino, Moscow region. Animals were housed under standard conditions (12:12 hr light/dark regime starting at 8:00 am, at a constant temperature of 22+/-2°C, with food in pellets and water available *ad libitum*). Mice were weaned at three weeks of age and housed in groups of 8-10 in standard plastic cages. Experiments were performed with 10-12 week old animals. All procedures were in compliance with the European Communities Council Directive 210/63/EU on September 22, 2010. The study was approved by Scientific Council N 9 of the Institute of Cytology and Genetics SD RAS of March, 24, 2010, N 613 (Novosibirsk).

### Generation of alternative social behaviors under agonistic interactions in male mice

Prolonged negative and positive social experience, social defeats and wins, in male mice were induced by daily agonistic interactions [Kudryavtseva, 1991; Kudryavtseva *et al.* 2014]. Pairs of weight-matched animals were each placed in a cage (14 x 28 x 10 cm) bisected by a perforated transparent partition allowing the animals to see, hear and smell each other, but preventing physical contact. The animals were left undisturbed for two or three days to adapt to new housing conditions and sensory contact before they were exposed to encounters. Every afternoon (14:00-17:00 p.m. local time), the cage lid was replaced by a transparent one, and 5 min later (the period necessary for individuals’ activation), the partition was removed for 10 minutes to encourage agonistic interactions. The superiority of one of the mice was firmly established within two or three encounters with the same opponent. The superior mouse would be attacking, biting and chasing another, who would be displaying only defensive behavior (sideways postures, upright postures, withdrawal, lying on the back or freezing). As a rule, aggressive interactions between males are discontinued by lowering the partition if the sustained attacks has lasted 3 min (in some cases less) thereby preventing the damage of defeated mice. Each defeated mouse (defeater, loser) was exposed to the same winner for three days, while afterwards each loser was placed, once a day after the fight, in an unfamiliar cage with an unfamiliar winner behind the partition. Each winning mouse (winners, aggressive mice) remained in its original cage. This procedure was performed once a day for 20 days and yielded an equal number of the winners and losers.

Three groups of animals were used: 1) Controls – mice without a consecutive experience of agonistic interactions; 2) Losers – chronically defeated mice; 3) Winners – chronically aggressive mice. The losers and winners with the most expressed behavioral phenotypes were selected for the transcriptome analysis. The winners and losers, 24 hours after the last agonistic interaction, and the control animals were simultaneously decapitated. The brain regions were dissected by one experimenter according to the map presented in the Allen Mouse Brain Atlas (http://mouse.brain-map.org/static/atlas). All biological samples were placed in RNAlater solution (Life Technologies, USA) and were stored at −70° C until sequencing.

The brain regions were selected for the analysis based on their functions and localization of neurons of neurotransmitter systems. These are as follows: the midbrain raphe nuclei, a multifunctional brain region, which contains the bodies of serotonergic neurons; the ventral tegmental area (VTA), which contains the bodies of dopaminergic neurons, is widely implicated in natural reward circuitry of the brain and is important for cognition, motivation, drug addiction, and emotions relating to several psychiatric disorders; the striatum, which is responsible for the regulation of motor activity and stereotypical behaviors and is also potentially involved in a variety of cognitive processes; the hippocampus, which belongs to the limbic system, is essential for memory consolidation and storage, and plays important roles in neurogenesis and emotional mechanisms; and the hypothalamus, which regulates the stress reaction and many other physiological processes.

### RNA-Seq

We used the methods described earlier [Smagin *et al.* 2016a; Smagin *et al*. 2016b]. The collected samples were sequenced at JSC Genoanalytica (www.genoanalytica.ru, Moscow, Russia), and the mRNA was extracted using a Dynabeads mRNA Purification Kit (Ambion, Thermo Fisher Scientific, Waltham, MA, USA). cDNA libraries were constructed using the NEBNext mRNA Library PrepReagent Set for Illumina (New England Biolabs, Ipswich, MA USA) following the manufacturer’s protocol and were subjected to Illumina sequencing. More than 20 million reads were obtained for each sample. The resulting “fastq” format files were used to align all reads to the GRCm38.p3 reference genome using the TopHat aligner [Trapnel *et al.* 2013]. DAVID Bioinformatics Resources 6.7 (http://david.abcc.ncifcrf.gov) was used for the description of differentially expressed gene ontology. The Cufflinks program was used to estimate the gene expression levels in FPKM units (fragments per kilobase of transcript per million mapped reads) units and subsequently identify the differentially expressed genes in the analyzed and control groups. Each brain area was considered separately for 3 vs 3 animals. Genes were considered differentially expressed at *P ≤* 0.05 and for multiplied comparisons q < 0.05.

We have previously conducted studies of gene expression in males in similar experiments using the RT-PCR method with larger number of samples for each compared experimental group, i.e., winners and losers (> 10 animals). The direction and extent of changes in expressions of the *Tph2, Slc6a4, Bdnf, Creb1*, and *Gapdh* genes in the midbrain raphe nuclei of males compared with the control produced by two methods, including RT-PCR [Smagin *et al.* 2013; Boyarskikh *et al.* 2013] and RNA-Seq [Kudryavtseva et al., 2017], are generally consistent. In order to cross-validate the our results obtained, we employed the unique resource from Barres Lab, Stanford University, USA [Zhang et al. 2014] and found highly concordant with our RNA-Seq data pool [Babenko et al. 2017]. As additional cross-validation may be considered positive correlation (r = 0.74, df = 42, *p* < 10E-4) between FPKM values of *Col\** family genes in our experiment and in the experiment of Kadakkuzha et al. [2015] in the hippocampus of C57BL/6J mice. These findings suggest that transcriptome analyses of the data provided by the ZAO Genoanalitika (http://genoanalytica.ru, Moscow) have been verified, and this method reflects the actual processes that occur in the brain under our experimental paradigm.

The Human Gene Database (http://www.genecards.org/); Online Mendelian Inheritance in Man database (OMIM, http://omim.org/); Human disease database (MalaCards, http://www.malacards.org) were used for the description and analysis of the data obtained.

### Statistical analysis

For the transcriptomic data, a Principal Components (PC) analysis was conducted using the XLStat software package (www.xlstat.com). It was based on a Pearson correlation metric calculated on the FPKM value profiles of 49 analyzed genes. Agglomerative Hierarchical Clustering (AHC) was performed on the same data with the XLStat software package. We also used a Pearson correlation as a similarity metric for the AHC analysis. The agglomeration method comprised an unweighted pair-group average.

## Results and discussion

Data of the RNA-Seq profiling in FPKM units for 49 collagen genes, including *Colgalt1* and *Colgalt2* (collagen beta(1-O)galactosyltransferase 1 and 2); *Colq* (collagen-like tail subunit of asymmetric acetylcholinesterase; *Adipoq* (adiponectin, C1Q and collagen domain containing, mRNA). Additionally, *Ccbe1* (collagen and calcium binding EGF domains 1, mRNA), as well as procollagen genes (*P4ha1, P4ha2, P4ha3, Pcolce, Pcolce2, Plod1, Plod2, Plod3*) were considered.

Analyzing whole transcriptome data in the hypothalamus, midbrain raphe nuclei, hippocampus, ventral tegmental area (VTA), and striatum, we found similar and different changes in the expression of *Col\** family genes in brain regions of male mice with alternative social experience of agonistic interactions (Table 1).

**Table 1.** Differentially expressed *Col*\* family genes in the brain regions of male mice with repeated experience of agonistic interactions (P < 0.05)

|  |  |  |  |  |  |
| --- | --- | --- | --- | --- | --- |
|  | MRN | HPC | VTA | STR | HPT |
| **Winners** |  |  |  |  |  |
| Genes, N | 6 | 5 | 12 | 5 | 19 |
| Upregulation | 2 | 2 | 0 | 5 | 19 |
| Downregulation | 4 | 3 | 11 | 0 | 0 |
| **Losers** |  |  |  |  |  |
| Genes, N | 8 | 15 | 9 | 20 | 19 |
| Upregulation | 4 | 14 | 2 | 17 | 16 |
| Downregulation | 4 | 1 | 7 | 3 | 3 |
| Note: MRN – midbrain raphe nuclei; HPC – hippocampus; VTA – ventral tegmental area; STR – striatum; HPT – hypothalamus. Winners – aggressive mice; Losers - defeated mice |  |  |  |  |  |

The most of *Col\** genes that changed their expression were found in the hypothalamus (19 genes in both groups), hippocampus and striatum (15 and 20 genes, respectively, in the losers). The minority of *Col\** differentially expressed genes were found in the midbrain raphe nuclei (6 and 8 genes in the winners and losers, respectively), in the hippocampus and striatum (for both 5 genes) in the winners and in the VTA (12 and 9 genes in the winners and losers, respectively). The most of differentially expressed genes of *Col\** family in the hypothalamus of the winners and losers as well as in the striatum and hippocampus of the losers were upregulated. In the VTA these genes were downregulated in mice of both groups. (Full list of differentially expressed genes in the winners and losers in different brain regions are given in the Section Results, and in Supplement: Table 1 and additional statistics for differentially expressed *Col\** genes in brain regions.

**In the hypothalamus** differentially expressed *Col4a2, Col5a3, Col9a2, Col9a3, Col11a2, Col16a1, Col18a1, Col22a1, Col23a1, Col24a1, Col27a1* and *Plod3* genes were upregulated in both social groups (Fig 1; Supplement, Table 1). Changes of the *Col4a1, Col6a3, Col11a1, Col15a1, Col26a1, and Plod1* expression were upregulated and *Col3a1, Col13a1* and *Plod2* genes were downregulated specifically in the losers.

**Figure 1.**
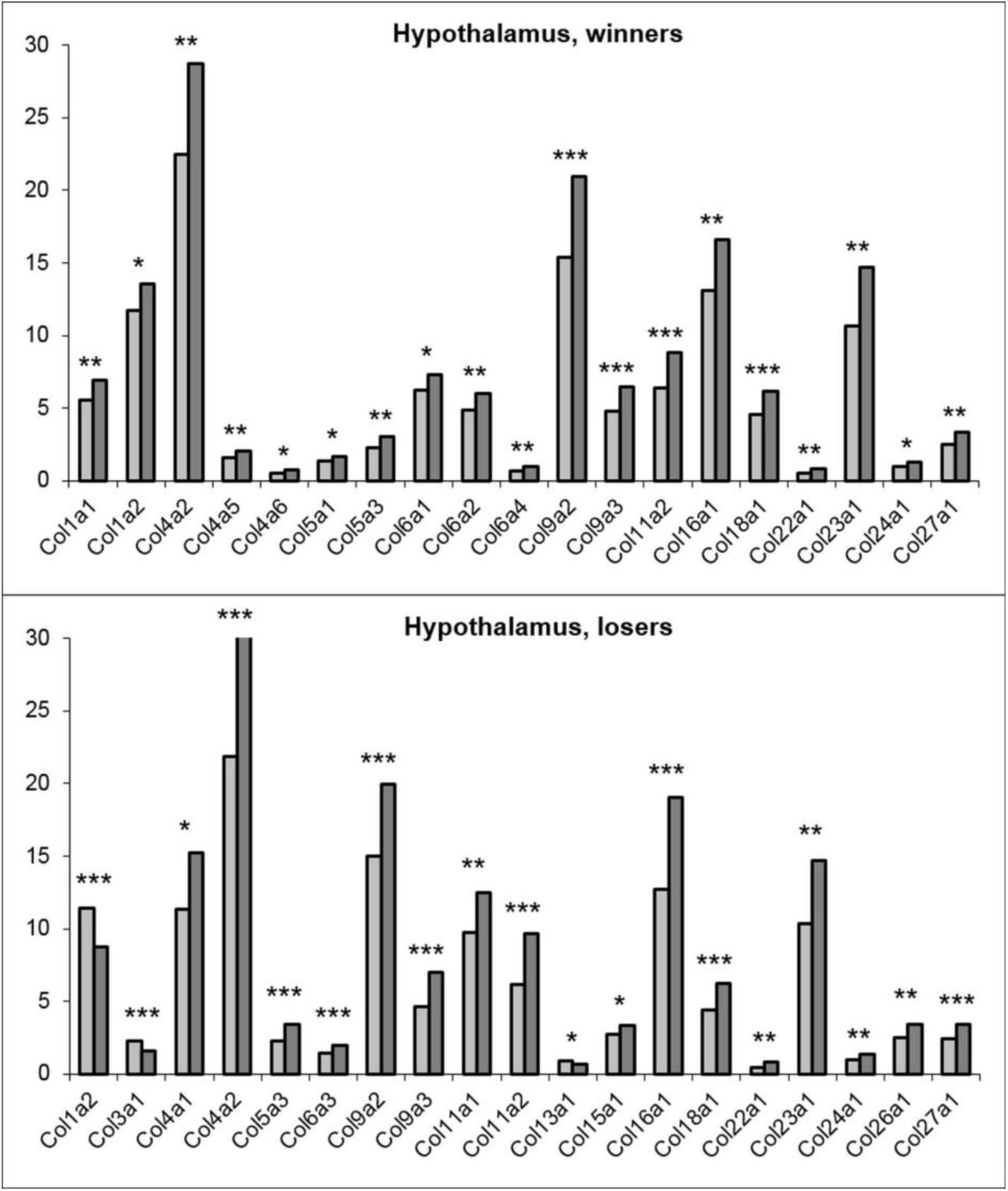
The differentially expressed collagen *Col\** family genes in the hypothalamus of mice with experience of repeated agonistic interactions. The Cufflinks program was used to estimate the gene expression levels in FPKM units. The levels of the *Col\** gene expression are presented in the control (left columns) and experimental mice (right columns) at the statistical significance * *P* < 0.05; \*\**P* < 0.01; \*\*\**P* < 0.001. Additional statistics was shown in Supplement.

The expression of the *Col1a1, Col4a5, Col4a6, Col5a1, Col6a1, Col6a2, Col6a4* and *Pcolce* genes was specifically upregulated in the winners. Expression of the *Col4a2, Col9a2*, and *Col16a1* and its changes were maximal in the both social groups. The *Col1a2* gene was upregulated in the winners and downregulated in the losers.

The upregulation of most *Col\** genes in the hypothalamus of the winners and losers may be the response to the chronic social stress inducing development of anxiety in both participants of social conflicts [Kudryavtseva & Avgustinovich 1998; Kudryavtseva *et al.* 2002]. Earlier we found similar upregulation of most amount of ribosomal (*Rpl*\* and *Rps*\* families) and mitochondrial ribosomal (*Mrpl*\* and *Mrps*\* families) genes in the hypothalamus of the winners and losers [Smagin *et al.* 2016a; Smagin *et al.* 2016b]. It’s therefore natural to suppose the coexpression or joint expression elevation of numerous genes developing under chronic social stress of agonistic interactions.

The *Col1a2, Col3a1* and *Col13a1* genes decreased their expression in the hypothalamus of the losers. According GeneCards database the *Col1a2* and *Col3a1* genes are associated with each other, and both participate in regulation of vascular system. Function of *Col3a1* gene product is not known, however, it has been detected at low levels in all connective tissue-producing cells. Unlike most of the collagens, which are secreted into the ECM, *Col3a1* gene protein has been localized to the plasma membrane, involved in cell-matrix and cell-cell adhesion interactions that are required for normal development. In the winners the *Col1a2* gene was upregulated oppositely to the losers. It may be supposed that this gene may be responsible for specific expression of *Col\** genes in the hypothalamus in every social group. This allows assuming differences in the state of the ECM in the hypothalamus of animals with alternative social experience.

Little is known about the specific role of collagen genes in brain regions under functional uploads. It was previously shown that mRNA expression for the *Col1a1* and *Col3a1* genes was distinctly observed in pituitary glands of rats in cells located around capillaries in the gland and were supposed to be the main components of the ECM [Fujiwara *et al.* 2010]. Interestingly, in the hypothalamus of Djungarian hamsters, transcriptome analysis revealed upregulation of the *Col18a1* and *Col5a3* similar with our results, and increase of the *Col17a1, Col20a1* and *Col26a1* gene expression, which were specific for torpor in comparison with normothermic hamsters [Cubuk *et al.* 2017]. The preliminary hypothesis from this part of data may be the recognition of involvement *Col\** genes into natural regulation, for example, of physiological adaptation under repeated stress in mice, winter torpor in hamsters, or CNS pathologies associated with collagens during neurodevelopment [Ali *et al.* 1998; Cheng *et al.* 2009; Lepelletier, *et al.* 2017].

**In the midbrain raphe nuclei** (Fig 2; Supplement, Table 1) expression of most genes therein was the lowest in comparison with other regions (< 5.0 FPKM units). Nevertheless, similar with the hypothalamus, expression of the *Col9a2* gene was upregulated in comparison with other genes. The *Col6a2, Col15a1, Col24a1,* and *Col25a1* genes were downregulated and the *Col6a3* and *Col9a2* genes were upregulated similarly in both social groups. Specific alteration was upregulation of the *Col6a1, Ccbe1* and *Colq* genes expression in the losers. Upregulation of the *Plod2* and *Plod3* genes and downregulation of the *Plod1* gene expression were specific for the winners (Supplement, Table 1).

**Figure 2.**
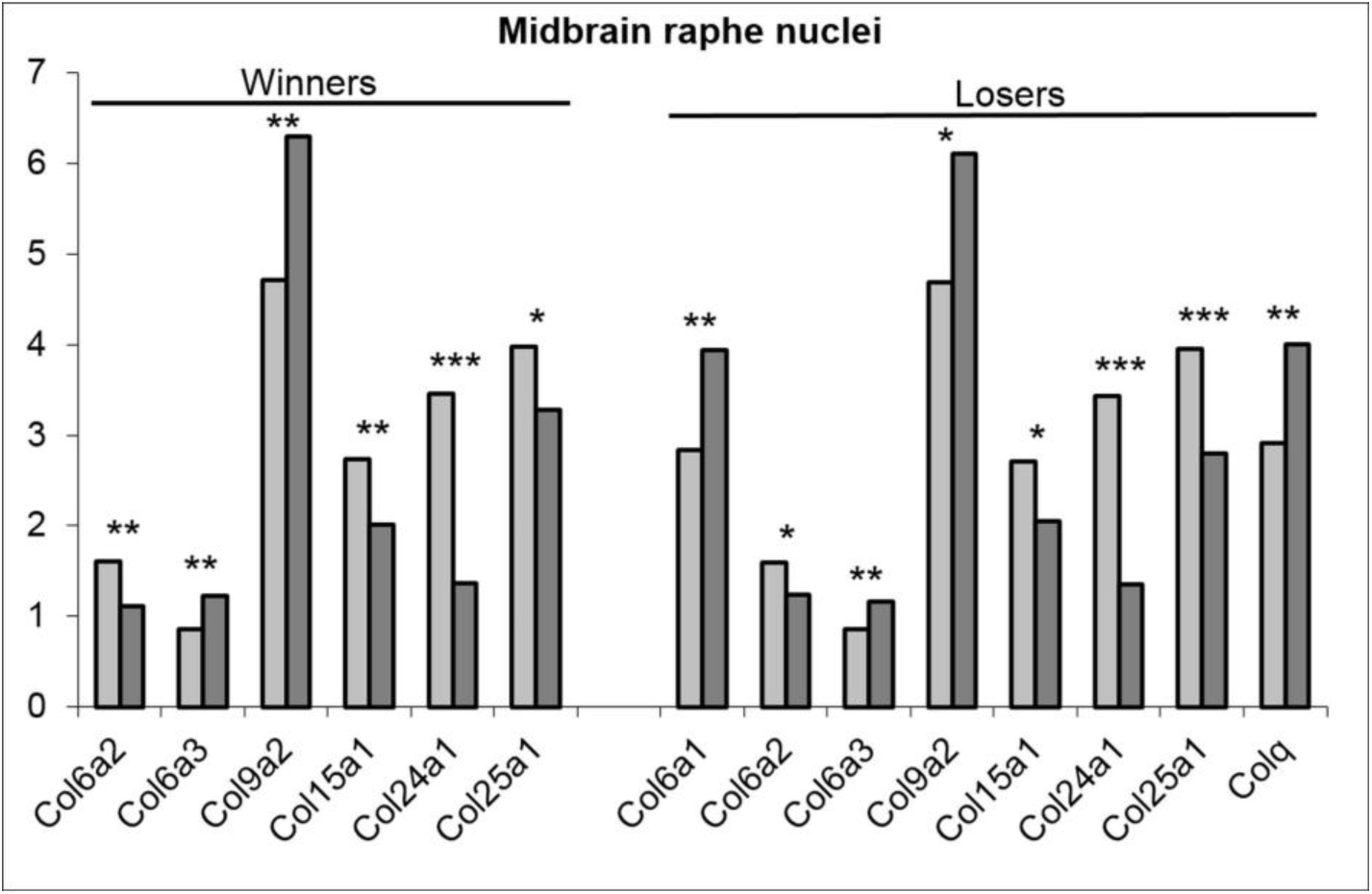
The differentially expressed collagen *Col\** family genes in the midbrain raphe nuclei of mice with agonistic interactions. The Cufflinks program was used to estimate the gene expression levels in FPKM units. The levels of the *Col* gene expression are presented in the control (left columns) and experimental mice (right columns) at the statistical significance * *P* < 0.05; \*\**P* < 0.01; \*\*\**P* < 0.001. Additional statistics was shown in Supplement.

The midbrain raphe nuclei contain the bodies of serotonergic neurons which are involved in the regulation of many physiological, behavioral, and emotional processes. Repeated experience of aggression and defeats is accompanied by the decrease of serotonergic activity [Kudryavtseva 2006; Avgustinovich *et al.* 2004], which is accompanied by decreased expression of serotonergic genes - *Tph2, Maoa, Slc6a4, Htr’s* [Boyarskikh *et al*. 2013; Smagin *et al*. 2013; Kudryavtseva *et al*. 2017] in this brain region.

It has been shown, that stimulation of serotonin production may enhance the synthesis of some collagen proteins in human mesangial cells [Kasho *et al*., 1998]. On the other hands, serotonin-dependent decrease in collagen mRNA was accompanied by decreased transcription serotonin, downregulating the gene for type I collagen and other the ECM proteins in myometrial cells [Passaretti *et al*. 1996]. Thus, we can suggest that decrease of some *Col\** gene expression in the midbrain raphe nuclei of the winners and losers may be associated with decreased serotonergic activity.

**In the VTA** (Fig 3, Supplement, Table 1) all genes excepting the *Col25a1* and *Colgalt2* genes in the losers were downregulated in both groups with different social experience: the *Col1a2, Col5a2, Col5a3, Col11a2, Col24a1,* and *Colq* genes lowed their expression similarly in the winners and losers under agonistic interactions. The *Col1a1, Col4a1, Col9a2, Col9a3, Col16a1*, and *Col23a1* genes specifically in the winners and the *Col27a1* specifically in the losers were downregulated. Maximal overall expression manifested *Col4a1, Col11a2, and Col16a1* genes in the winners and the *Col11a2* gene in the losers.

**Figure 3.**
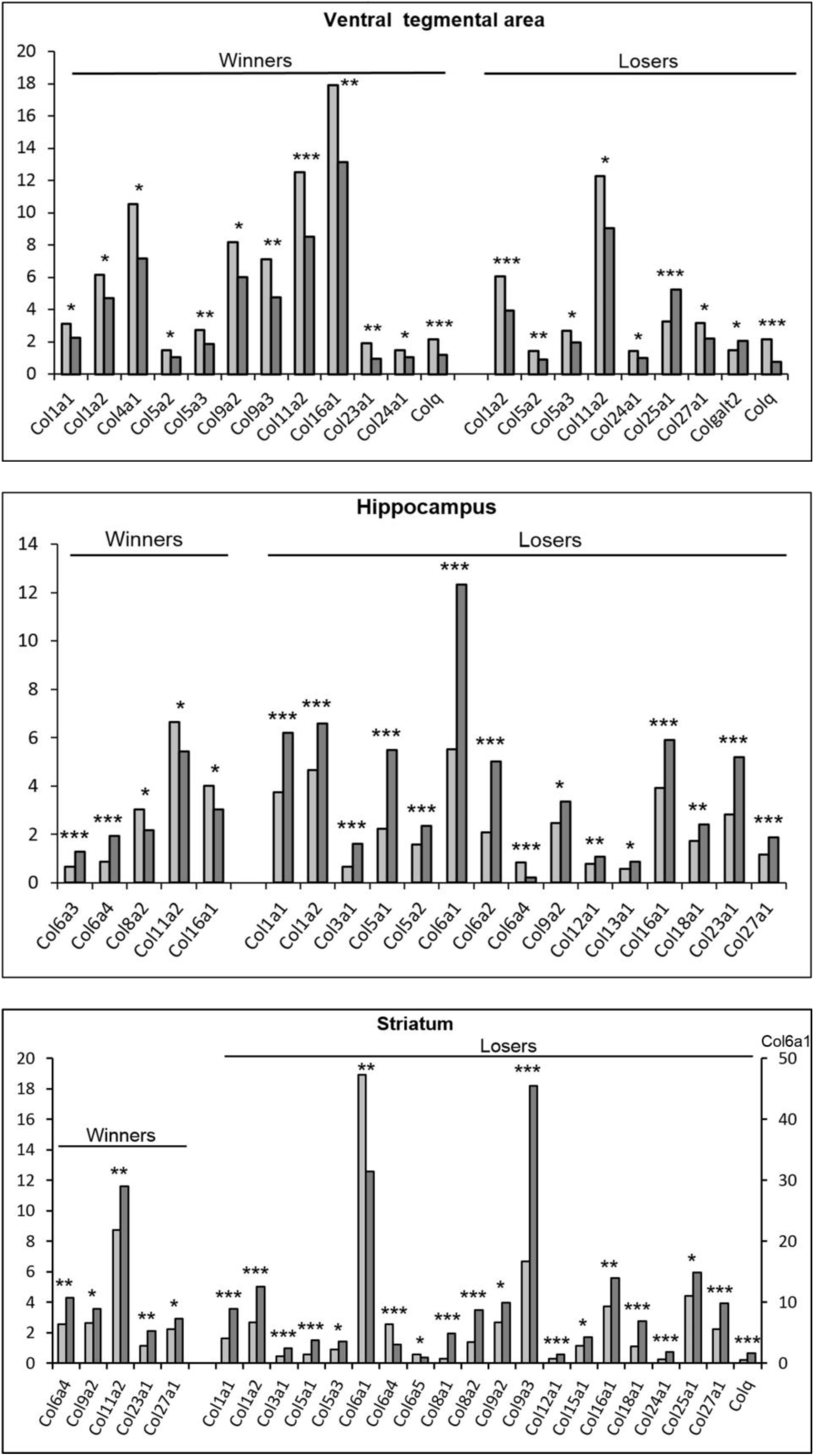
The differentially expressed collagen *Col\** family genes in the ventral tegmental area, hippocampus and striatum of mice with experience of repeated agonistic interactions. The Cufflinks program was used to estimate the gene expression levels in FPKM units. The levels of the *Col* gene expression are presented in the control (left columns) and experimental mice (right columns) at the statistical significance * *P* < 0.05; \*\**P* < 0.01; \*\*\**P* < 0.001. Additional statistics was shown in Supplement.

Comparing the changes of *Col\** gene expression in mice under agonistic interactions in the VTA and midbrain raphe nuclei, we noticed that only the *Col24a1* gene similarly decreased expression in both brain regions in both social groups, while other genes which changed their expression were different. The identical genes, for example, the *Col9a2* in the winners and the *Col25a1* and *Colq* gene in the losers changed their expression oppositely in the midbrain raphe nuclei and VTA. That may indicate different processes in the ECM in brain regions. It’s worth to notice that VTA and midbrain raphe nuclei contain pericarions of neurons. Serotonergic neurons are predominant in the midbrain raphe nuclei. The VTA contains 50-70% dopaminergic neurons [Margolis *et al*. 2006; Nair–Roberts *et al*. 2008; Yamaguch *et al*. 2007], about 30% of GABAergic neurons [Bourdy & Barrot 2012] and ∼2-3% of glutamatergic neurons [Margolis *et al*. 2006; Nair–Roberts *et al*. 2008]. We can assume that the type of collagen genes involved in the effects of agonistic interactions may be due to the difference in neurotransmitter systems involved in this process in various brain structures

**In the hippocampus** (Fig 3, Supplement, Table 1) expression of the *Col1a1, Col1a2, Col3a1, Col5a1, Col5a2, Col6a1, Col6a2, Col9a2, Col12a1, Col13a1, Col18a1, Col23a1, Col27a1* genes were specifically upregulated in the losers and the *Col6a3* gene was specifically upregulated in the winners. Expression of *Col11a2* and *Col8a2* genes was downregulated in the winners. Interestingly, expression of the *Col6a4* gene was increased in the winners and was decreased in the losers. The *Col16a1* gene changed the expression oppositely in these social groups. Again, similar with VTA, we can suppose different changes in the ECM in male mice with alternative social experience.

Interestingly, all of the ribosomal and mitoribosomal genes were upregulated in the winners in this region [Smagin *et al.* 2016b] and most of them were downregulated in the losers [Smagin *et al.* 2016a]. These dynamics in changes of expression of *Rp\** genes we connected with different cell proliferation in this brain region. Repeated aggression has been shown to be accompanied by an increase in proliferation of neuronal progenitors and by production of new neurons in the dentate gyrus of the hippocampus [Smagin *et al.* 2015]. At the same time, other authors [Ferragud *et al.* 2010; Lagace *et al.* 2010; Van Bokhoven *et al.* 2011] have demonstrated the decreased cell proliferation in this brain region in defeated mice. It was suggested that the change in processes of neurogenesis is the consequence of ribosomal dysfunction developing under chronic positive or negative social experience in our experimental paradigm. Now we can suppose association between the decreased neurogenesis, the downregulation of ribosomal gene expression and the upregulation of *Col\** genes expression in the losers and increased neurogenesis, upregulation of ribosomal gene expression and decreased *Col\** gene expression in the hippocampus in the winners. However, we cannot exclude the possibility of the opposite relationship: changes in neurogenesis may influence the ribosomal and collagen functions.

Association between neurogenesis and changes in *Col\** gene expression is indirectly confirmed by experiments [Kakoi *et al.* 2012]: administration for 4 weeks orally of the lower molecular weight peptides derived from collagen enhanced the hippocampal neurogenesis and exerted anxiety-related behavior in adult mice: the density of proliferating cells in subgranular zone showed a 1.2-fold increase. Moreover, there is a growing volume of papers underlining that hippocampal functioning and axon outgrowth are tightly connected with collagen genes activation [Xia et *al.* 2013; Carletti *et al.* 2016]. Our preliminary analysis of brain specific genes underlined vesicular glutamate transporter *Slc17a7* (VGLUT1) as strictly hippocampus specific highly expressed gene. In particular, we observed very intense expression rate of vesicle transporting gene *Slc17a7* specifically in the hippocampus samples of our mice (Figure 1). This gene was shown to be implicated in neuroendocrine response upon chronic stress experience [Myers et al. 2017]. Some data suggest a functional role of SLC17A7 in the hippocampal synaptic plasticity [Balschun *et al*. 2010; Li et al. 2011; Bogen et al. 2009], though there is no direct evidence of its activity and synapse plasticity [Fung et al., 2011]. The expression pattern of *Slc17a7* gene in our data was highly correlated with at least three collagen genes: *Col19a1* (*r* = 0.88; *df* = 43; *P* < 1E-6), *Col5a1* (*r=* 0.77; *df* = 43; *P* < 1E-5), *Col4a1* (*r* = 0.4; *df* = 43). Notably, all of these proteins express specifically in neuronal cells [Zhang *et al.* 2014]. It underscores the ultimate importance of collagen families of the reported previously axon outgrowth and synapse activation in hippocampus specifically upon the stress experience [Hayashi, 2015]. In particular, the neurons will grow their neuritis along the collagen matrix since they maintain receptors on their plasma membrane adhesive to the collagen. The implications of collagen deviated expression may signal on specific events in synapse and axon ramifications/functions. One of other points worth mentioning about neuron-specific collagen activities is that the above mentioned *Col5a1* expression rate is manifested as hippocampus specific, at the same time highly elevated in the hippocampus of depressive mice in our data. This elevation is highly significant and consistent. We should note that such consistency in a group - specific expression pattern is quite rarely observed in ours and other data.

**In the striatum (**Fig 3, Supplement, Table 1) the *Col6a4, Col9a2*, and *Col27a1* genes were upregulated in both social groups. The upregulation of gene expression specific for the winners was shown for the *Col11a2* and *Col23a1* genes, and specific for losers – for the *Col1a1, Col1a2, Col3a1, Col5a1, Col5a3, Col8a1, Col8a2, Col9a3, Col12a1, Col15a1, Col16a1, Col18a1, Col24a1, Col25a1, Colq, Pcolce2, Pcolce, Plod1,* and *Cthrc1* genes. Decreased expression was revealed for the *Col6a1, Col6a4* and *Col6a5* genes in the losers. Similarly with the hippocampus the *Col6a4* gene was upregulated in the winners and downregulated in the losers. Highest expression was shown for the *Col6a1* gene in the losers.

The striatum is responsible for the modulation of movement pathways and is potentially involved in a variety of other cognitive processes responsible for regulation of motor activity and stereotypic behaviors. The winners demonstrate hyperactivity and enhanced locomotion in behavioral tests and stereotypical behaviors, *etc*. [Kudryavtseva 2006; Kudryavtseva *et al.* 2014; Vishnivetskaya *et al.* 2013]. In the losers with low locomotor and exploratory activity, helplessness, and immobility behavior in any situation, the total number of differentially expressed genes was 4 times more than in the winners, and the majority of these genes in this brain region were upregulated [Smagin *et al.* 2017]. In spite of differences in the locomotor behavior in the winners and losers, the majority of differentially expressed genes were upregulated in both social groups.

### Principal components analysis (PC) of Col* gene expression in the RNA-Seq data

To assess the degree of brain region-specific expression of genes of interest, we performed a Principal Components analysis based on the co-variation of 49 genes using the expression profiles of 45 samples, which comprised RNA-Seq FPKM data for 5 brain regions of 9 mice. Ovals correspond to brain regions. We identified compact clustering of the hypothalamus, striatum and hippocampus samples based on gene expression profiles (Fig 4, encircled), whereas the midbrain raphe nuclei and ventral tegmental area were merged.

**Figure 4.**
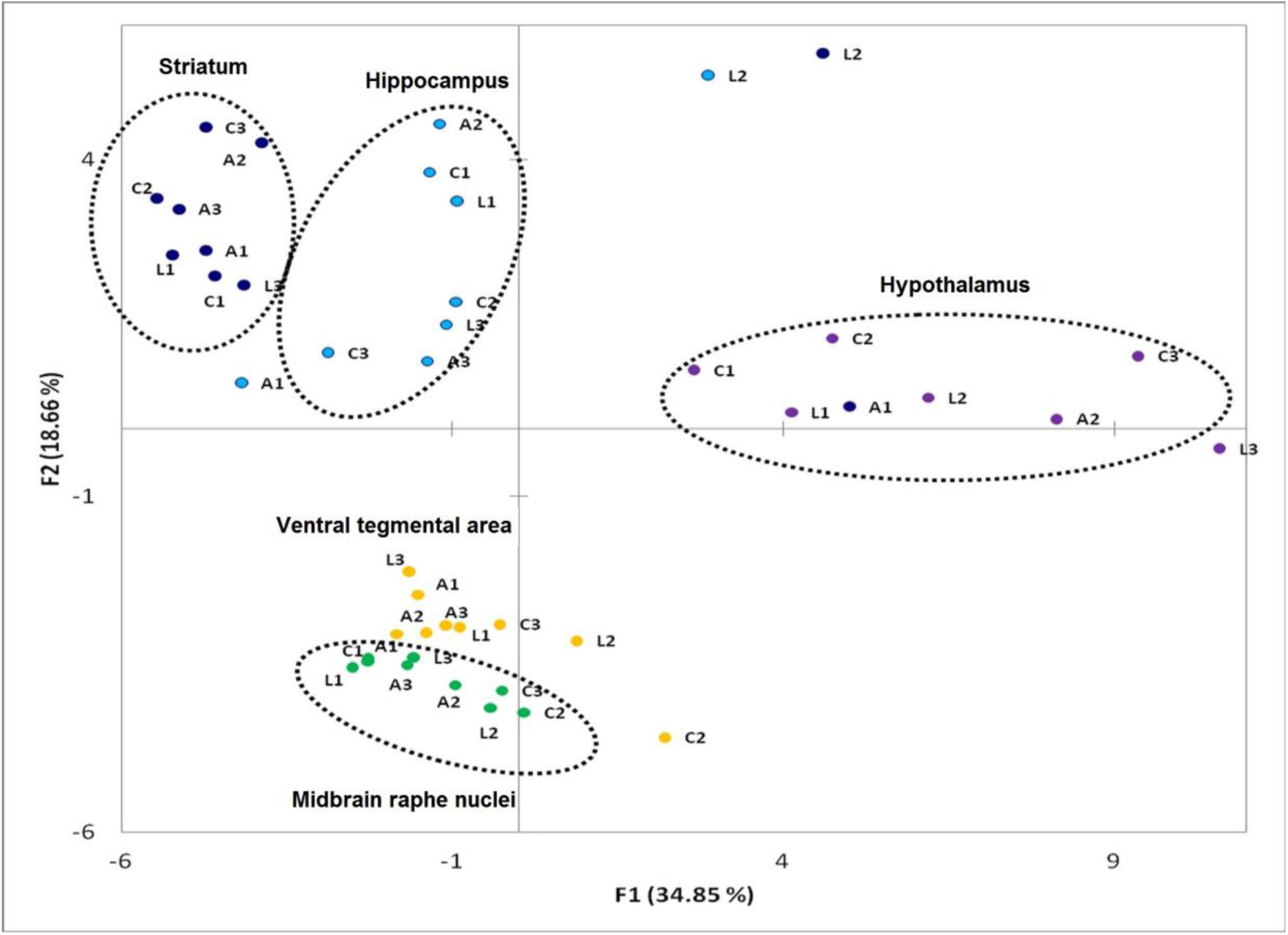
Principal component analysis plot based on co variation of 49 *Col\** family genes using the expression profiles of 45 samples, which comprised RNA-Seq FPKM data for 5 brain regions of 9 mice. Ovals correspond to brain regions. C1, C2, C3 – control; A1, A2, A3 – aggressive mice; L1, L2, L3 – defeated mice, losers. Distinct clustering of three brain regions occurred, whereas the midbrain raphe nuclei and ventral tegmental area were not distinct.

### Agglomerative hierarchical clustering (AHC) of Col* genes coding transcripts and identification of outliers

We applied AHC to the initial sample of 49 genes, based on their expression profiles (Fig. 5) presents the dendrogram of gene clustering. The similarity ordinate corresponds to the Pearson correlation coefficient (df = 44). An AHC analysis elucidates closely correlated genes, which thus strengthens the confidence of concordant differential expressions of a gene pair based on their highly correlated expression profiles across 45 samples. Thus, we reaffirm the significant differential expression given correlated pairs of genes and independent tests using the Cufflinks program. The overlapping or specific *Col\** genes in different brain areas of the winners and losers are presented in Supplement, Table 1.

**Figure 5.**
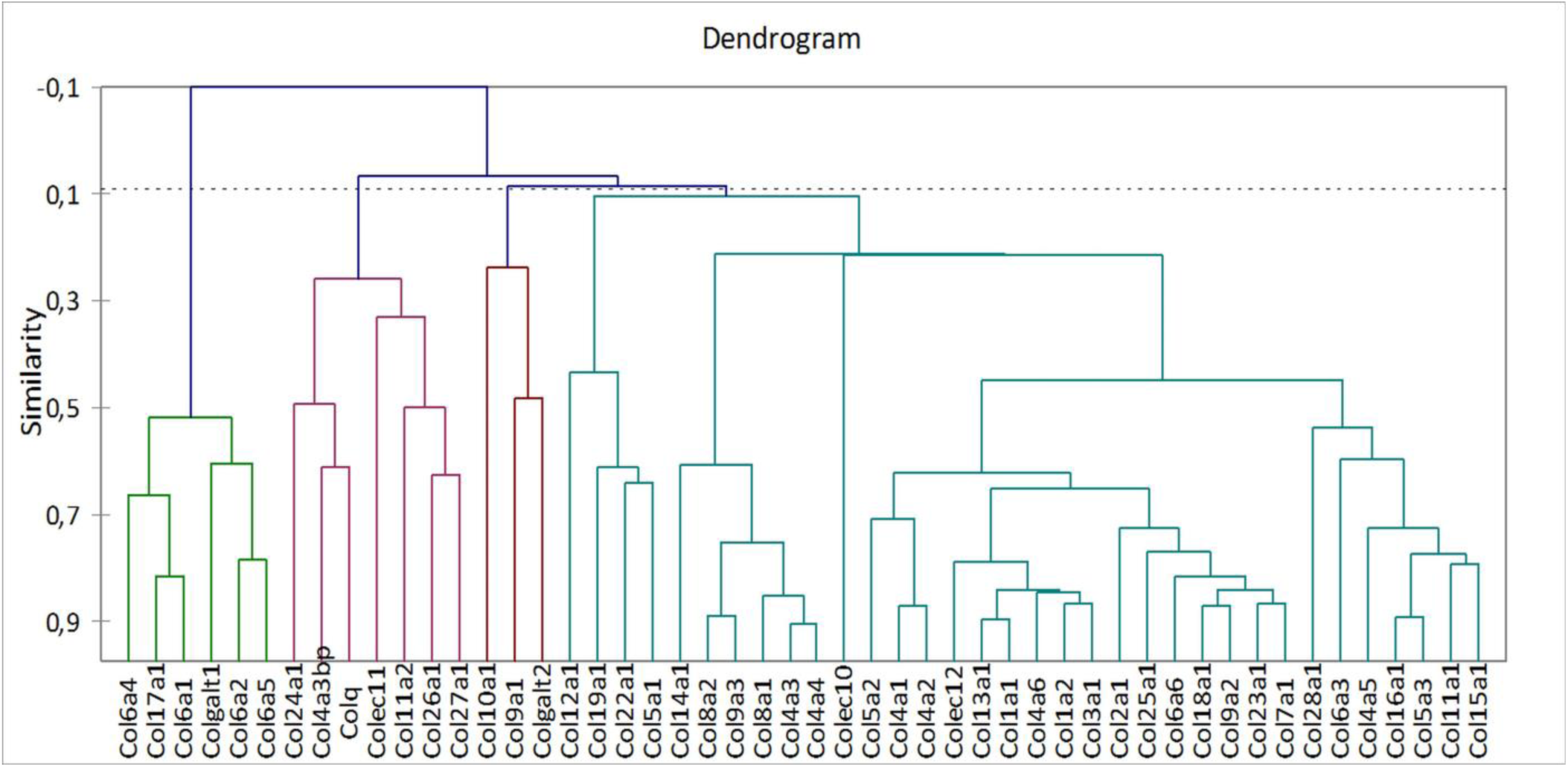
Agglomerative hierarchical clustering of *Col\** coding transcripts and identification of outliers. We applied agglomerative hierarchical clustering to the initial sample of 49 genes across 45 samples, which comprised RNA-Seq FPKM data for 5 brain regions of 9 mice based on their expression profiles. Figure presents the dendrogram of gene clustering. The similarity ordinate corresponds to the Pearson correlation coefficient (df=44).

## General discussion

We should underline principal points in dealing with collagen genes. We found the effect for brain regions in the winners and losers, and we were able to see consistent elevations and downturns of expression rate for particular groups of collagen proteins across brain regions that were elucidated by PC analysis underlining their brain region specific expression profile.

In the hypothalamus, striatum and hippocampus the most of *Col\** genes were upregulated, and the amount of differentially expressed genes in the striatum and hippocampus was significantly more in the losers than in the winners. In the midbrain raphe nuclei and VTA expression of the most of *Col\** genes were downregulated. In has been shown that the direction of changes in expression of some *Col*\* genes depended also on social experience of mice: some genes changed their expression specific for the winners and for the losers.

We can assume the dysfunction of the ECM due to aberrant *Col\** gene expression. We can suppose also that changed neurotransmitter activity involved in the regulation of pathological states primarily creates the conditions for development of aberrant functioning of the ECM which, as consequence, produces changes in works of many metabolic genes including ribosomal and mitoribosomal genes shown earlier for brain regions [Smagin *et al.* 2016a; Smagin *et al.* 2016b].

It was proposed [Kerrisk 2014; Bonneh-Barkay & Wiley 2009] that collagens can serve many functions, such as mediating cell adhesion, segregating tissues from one another, and regulating intercellular connection, transmitting signals to cell surface adhesion receptors, and participating in the regulation of synaptic plasticity. Changes in physiological conditions can trigger the rapid and local growth factor-mediated activation of cellular functions without additional synthesis. The stiffness and elasticity of the ECM, as supposed, have important implications in cell migration, gene expression, and differentiation [Engler *et al.* 2006; Frantz *et al.* 2010; Wang *et al.* 2007]. The precise expression pattern of collagen genes depends on a balance of positive and negative transcription factors, proteins that control the synthesis of mRNA from the specific gene [Okazaki & Sandell 2004]. Authors suppose that during development or the disease, the specific genes must be expressed in order to make or repair appropriate ECM. In any case, there are dynamic interrelations between the ECM influences on gene expression via transmembrane proteins as was earlier supposed by previous hypothesis [Slavkin 1982; Bissell & Barcellos-Hoff 1987]. In turn, changes in expression of numerous genes may impact cell interactions with other cells, creating novel biogenic environment, which may be reason of aberrant *Col\** gene expression.

The majority of collagen disorders are associated with neurological abnormalities including seizures and myoclonus, psychomotor retardation, spasticity, motor neuron disease, weakness, chronic fatigue, and endocrine abnormalities *etc*. Aggressive and defeated mice have also been shown to demonstrate disturbances in motor activity and neurological symptoms [Kudryavtseva 2006; Kudryavtseva *et al.* 2014; Vishnivetskaya *et al.* 2013; Smagin *et al.* 2018]. We suggest that collagen abnormalities in male mice with alternative social experience can be considered as development of “functional collagenopathy” as opposed to a collagenopathy that is induced by mutations accompanied by alterations in the structure or function of collagen components.

At this stage of our research, it is impossible to elucidate the detailed succession of neurochemical and molecular events which occur as a result of changes in brain regulation under repeated agonistic interactions in male mice. However, it is clear that this process is starting with a change in social behaviors and, at a certain stage, launches a cascade of systemic changes in the whole brain, specific regions, metabolism, neurons, and the neurotransmitter systems, which leads to a change in the expression of numerous genes. We can assume that the aberrant expression of genes is a consequence but not a reason of different psychopathologies. The next stage of our work will be to find the key, major genes that run the entire chain of events from the changes in social behaviors to changes in the gene expression.

## Acknowledgments

This work was funded by Russian Science Foundation, grant no. 14-15-00063.

## Supplement

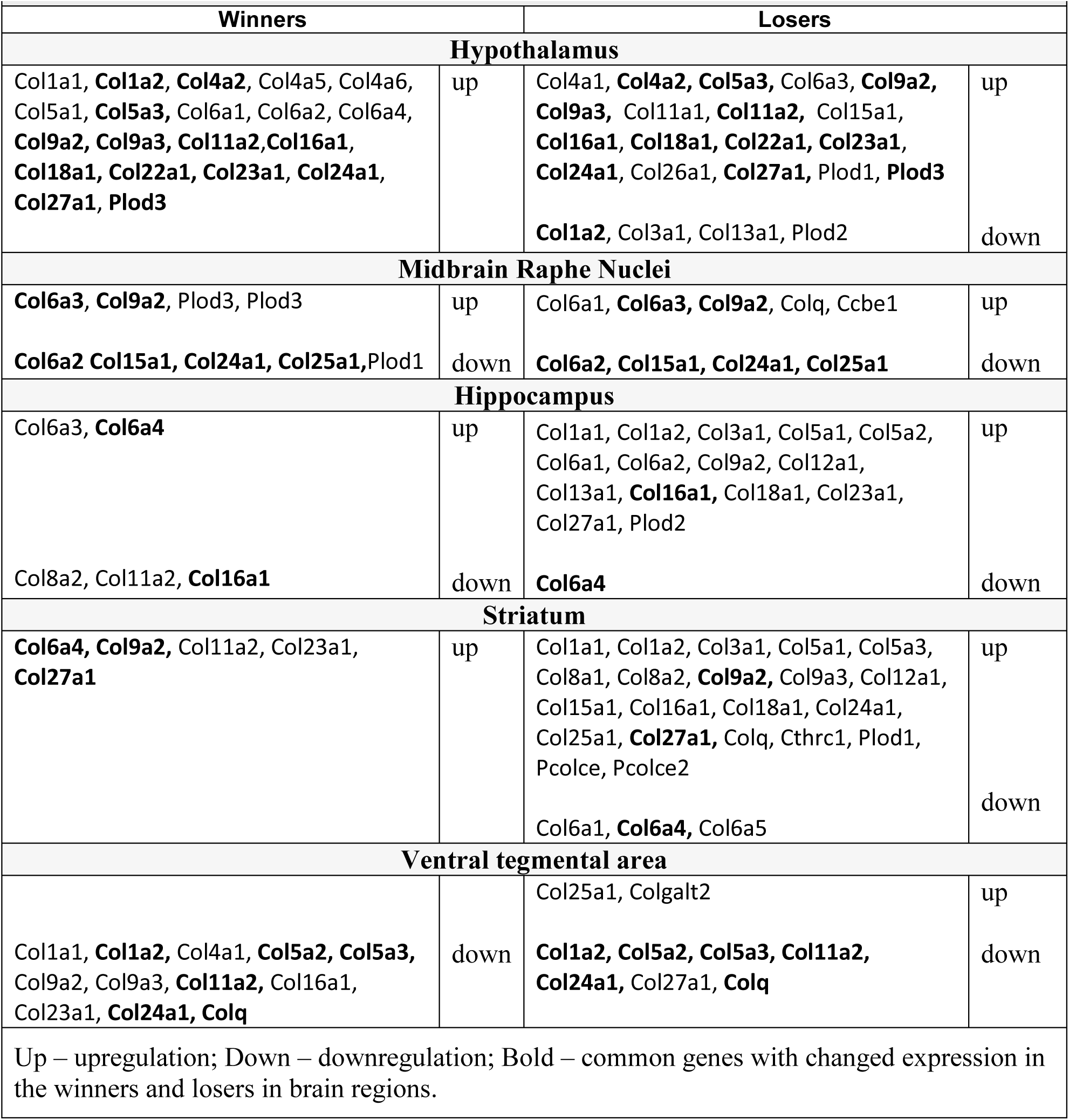
Differentially expressed Col* genes in brain regions of male mice with repeated experience of agonistic interactions.

| Table 1. Differentially expressed Col* genes in brain regions of male mice with repeated experience of agonistic interactions |  |  |  |
| --- | --- | --- | --- |
| Winners |  | Losers |  |
| Hypothalamus |  |  |  |
| Col1a1, **Col1a2**, **Col4a2**, Col4a5, Col4a6, Col5a1, **Col5a3**, Col6a1, Col6a2, Col6a4, **Col9a2**, **Col9a3**, **Col11a2**,**Col16a1**, **Col18a1**, **Col22a1**, **Col23a1**, **Col24a1**, **Col27a1**, Plod3 | up | Col4a1, **Col4a2**, **Col5a3**, Col6a3, **Col9a2**, **Col9a3**, Col11a1, **Col11a2**, Col15a1, **Col16a1**, **Col18a1**, **Col22a1**, **Col23a1**, **Col24a1**, Col26a1, **Col27a1**, Plod1, **Plod3** | up |
|  |  | **Col1a2**, Col3a1, Col13a1, Plod2 | down |
| Midbrain Raphe Nuclei |  |  |  |
| **Col6a3**, **Col9a2**, Plod3, Plod3 | up | Col6a1, **Col6a3**, **Col9a2**, Colq, Ccbe1 | up |
| **Col6a2** **Col15a1**, **Col24a1**, **Col25a1**,Plod1 | down | **Col6a2**, **Col15a1**, **Col24a1**, **Col25a1** | down |
| Hippocampus |  |  |  |
| Col6a3, **Col6a4** | up | Col1a1, Col1a2, Col3a1, Col5a1, Col5a2, Col6a1, Col6a2, Col9a2, Col12a1, Col13a1, **Col16a1**, Col18a1, Col23a1, Col27a1, Plod2 | up |
| Col8a2, Col11a2, **Col16a1** | down | **Col6a4** | down |
| Striatum |  |  |  |
| **Col6a4**, **Col9a2**, Col11a2, Col23a1, **Col27a1** | up | Col1a1, Col1a2, Col3a1, Col5a1, Col5a3, Col8a1, Col8a2, **Col9a2**, Col9a3, Col12a1, Col15a1, Col16a1, Col18a1, Col24a1, Col25a1, **Col27a1**, Colq, Cthrc1, Plod1, Pcolce, Pcolce2 | up |
|  |  | Col6a1, **Col6a4**, Col6a5 | down |
| Ventral tegmental area |  |  |  |
|  |  | Col25a1, Colgalt2 | up |
| Col1a1, **Col1a2**, Col4a1, **Col5a2**, **Col5a3**, Col9a2, Col9a3, **Col11a2**, Col16a1, Col23a1, **Col24a1**, **Colq** | down | **Col1a2**, **Col5a2**, **Col5a3**, **Col11a2**, **Col24a1**, Col27a1, **Colq** | down |
| Up – upregulation; Down – downregulation; Bold – common genes with changed expression in the winners and losers in brain regions. |  |  |  |

## Additional statistics for differentially expressed *Col\** genes in brain regions of male mice with repeated experience of agonistic interactions

**In the hypothalamus** the *Col\** genes increased their expression both in the winners and losers in comparison with the controls as for genes: *Col4a2* (*P* ≤ 0.003 and *P* < 0.0001; q < 0.001, respectively), *Col5a3* (*P* < 0.003; q < 0.047 and *P* < 0.0001; q < 0.002, respectively), *Col9a2* (*P* ≤ 0.0001; q < 0.003 and *P* < 0.0003; q < 0.004 respectively), *Col9a3* (*P* < 0.0004; q < 0.015 and *P* < 0.0001; q < 0.001, respectively), *Col11a2* (*P* < 0.0001; q ≤ 0.003, and *P* < 0.0001; q ≤ 0.001, respectively), *Col16a1* (*P* < 0.0031; q < 0.05 and *P* < 0.0001; q < 0.001, respectively), *Col18a1* (*P* < 0.0001; q ≤ 0.003 and *P* < 0.0001; q < 0.001, respectively), *Col22a1* (*P* < 0.0034 and *P* < 0.0041; q < 0.025, respectively*), Col23a1* (*P* < 0.0031 and *P* < 0.0099; q < 0.048, respectively); *Col24a1* (*P* < 0.011 and *P* < 0.0033; q < 0.0212, respectively), *Col27a1* (*P* < 0.0027; q < 0.05 and *P* < 0.0010; q < 0.009). Expression of *Col1a2* gene was upregulated in the winners (*P* < 0,043), and down regulated in the losers (*P* < 0.0004; q < 0.005). In comparison with the controls, the winners expression rate was increased in the *Col1a1* (*P* < 0.005), *Col4a5* (*P* < 0.005), *Col4a6* (*P* < 0.017), *Col5a1* (*P* < 0.017), *Col6a1* (*P* < 0.043), *Col6a2* (P < 0.008), *Col6a4* (P < 0.009) genes. In the losers, expression of the *Col4a1* (*P* < 0.024), *Col6a3* (*P* < 0.001, q < 0.001), *Col11a1* (*P* < 0.005, q < 0.029), *Col15a1* (*P* < 0.047) *Col26a1* (*P* < 0.009, q < 0.042) genes were increased and the *Col3a1* (*P* < 0.001, q < 0.006) and *Col13a1* (*P* < 0.045) genes were decreased. In the losers increased expression of the *Plod1* (*P* < 0.008; q < 0.041) and *Plod3 (P* < 0.0011; q < 0.010) genes as well as decreased expression of *Plod2* gene (*P* < 0.0002; q < 0.003) was found. In the winners increased expression of *Pcolce (P* < 0.006) and *Plod3* (*P* < 0.008) genes was found.

**In the midbrain raphe nuclei** in the winners and losers expression of the *Col6a3* (*P* < 0.002 and *P* < 0.009, respectively), *Col9a2* (*P* < 0.006 and *P* < 0.018, respectively), genes were increased and *Col6a2* (*P* < 0.005 and *P* < 0.049, respectively), *Col15a1* (*P* < 0.004 and *P* < 0.014, respectively), *Col24a1* (for both *P* < 0.0001; q < 0,005), *Col25a1* (*P* < 0.039 and *P* < 0.001, respectively) genes were decreased (Figure 2, Supplement Table 2, 4). Additionally in the losers expression of the *Col6a1* (*P* < 0.003), *Ccbe1 (P* <0.008) and *Colq* (*P* < 0.007) genes were upregulated. In the losers we found increased expression of protocollagen *Ccbe1* gene (*P* ≤ 0.007). In the winners decreased expression of *Plod1* gene (*P* < 0.025) and increased expression of *Plod2* (*P* < 0.0095) and *Plod3 (P* < 0.013) genes was found.

**In the VTA** of the winners and losers decreased expression of the *Col1a2* (*P* < 0.022 and *P* < 0.001, respectively), *Col5a2* (*P* < 0.014 and *P* < 0.002, respectively), *Col5a3* (*P* < 0.006 and *P* < 0.032, respectively), *Col11a2* (*P* < 0.001 and *P* < 0.012, respectively), *Col24a1*(for both *P* < 0.013), *Colq* (*P* < 0.0004 and *P* < 0.001; q < 0.007, respectively) genes were common for both social groups. The *Col1a1* (*P* < 0.016), *Col4a1* (*P* < 0.035), *Col9a2* (*P* < 0.015), *Col9a3* (*P* < 0.002), *Col16a1* (*P* < 0.007), *Col23a1* (*P* < 0.004) in the winners, and *Col27a1* (*P* < 0.013) in the losers specifically decreased their expression. In the losers *Col25a1* (*P* < 0.001; q < 0.021) and *Colgalt2* (*P* < 0.021) genes were upregulated.

**In the hippocampus** there are no genes that changed their expression in the same direction common for both social groups. However the increased expression of the *Col1a1* (*P* < 0.0001; q < 0,005), *Col1a2* (*P* < 0.0005; q < 0,029), *Col3a1* (*P* < 0.0001; q < 0,005), *Col5a1* (*P* < 0.0001; q < 0,005), *Col5a2* (*P* < 0.0005; q < 0,031), *Col6a1* (*P* < 0.0001; q < 0,005), *Col6a2* (*P* < 0.0001; q < 0,005), *Col9a2* (*P* < 0.015), *Col12a1* (*P* ≤ 0.007), *Col13a1* (*P* < 0.033), *Col16a1 P* < 0.0001; q < 0,0093), *Col18a1* (*P* < 0.0053), *Col23a1* (*P* < 0.001; q < 0,046), *Col27a1* (*P* ≤ 0.0003; q < 0,021) was revealed in the losers and *Col6a3* (*P* < 0.0001; q < 0,021) in the winners. The *Col6a4* gene were upregulated in the winners (*P* < 0.0004) and downregulated in the losers (*P* < 0.0001; q < 0,01). Oppositely, the *Col16a1* gene were downregulated in the winners (*P* < 0.014) and upregulated in the losers (*P* < 0.0001; q < 0,009). The *Col8a2* (*P* < 0.021) and *Col11a2* (*P* < 0.034) genes were specifically downregulated in the winners. Expression of the *Plod2* gene was decreased in the losers (*P* < 0.036).

**In the striatum** the *Col9a2* and *Col27a1* genes were upregulated both in the winners (*P* < 0.036 and *P* < 0.048) and losers (*P* < 0.024 and *P* < 0.001, q < 0.041). Specifically the *Col11a2* (*P* ≤ 0.01) and *Col23a1* (*P* < 0.002) genes in the winners and the *Col1a1* (*P* < 0.0001, q < 0.002), *Col1a2* (*P* < 0.0001, q <0.002), *Col3a1* (*P* < 0.0003, q < 0.015), *Col5a1* (*P* < 0.0001, q ≤ 0.004), *Col5a3 (P* < 0.027), *Col8a1* (*P* < 0.0001, q ≤ 0.003), *Col8a2* (*P* < 0.0001, q ≤ 0.003), *Col9a3* (*P* < 0.0001, q ≤ 0.003), *Col12a1* (*P* < 0.001, q < 0.04), *Col15a1* (*P* < 0.027), *Col16a1*(*P* < 0.009), *Col18a1* (*P* < 0.0001, q ≤ 0.003), *Col24a1* (*P* < 0.0001, q ≤ 0.003), *Col25a1* (*P* < 0.039), *Colq* (*P* < 0.0011) genes in the losers were upregulated and the *Col6a1* (*P* < 0.001, q ≤ 0.05) and the *Col6a5* (*P* < 0.025) were downregulated. The *Col6a4* gene was upregulated in the winners (*P* ≤ 0.002) and downregulated in the losers (*P* < 0.001, q ≤ 0.042). In the losers we found increased expression of genes *Cthrc1* (*P* < 0.0001, q ≤ 0.003), *Plod1* (*P* < 0.007), *Pcolce* (*P* < 0.0001, q ≤ 0.003), and *Pcolce2* (*P* < 0.004).

